# Apolipoprotein E4 disrupts the neuroprotective action of sortilin in neuronal lipid metabolism and endocannabinoid signaling

**DOI:** 10.1101/2020.01.12.903187

**Authors:** Antonino Asaro, Anne-Sophie Carlo-Spiewok, Anna R. Malik, Michael Rothe, Carola G. Schipke, Oliver Peters, Joerg Heeren, Thomas E. Willnow

**Author notes:** Correspondence to: Thomas E. Willnow, Max-Delbrueck-Center for Molecular Medicine, Robert-Roessle-Str. 10, D-13125 Berlin, Germany.

## Abstract

**INTRODUCTION:** ApoE is a carrier for brain lipids and the most important genetic risk factor for Alzheimer’s disease (AD). ApoE binds the receptor sortilin which mediates uptake of apoE-bound cargo into neurons. The significance of this uptake route for brain lipid homeostasis and AD risk seen with apoE4, but not apoE3, remains unresolved.

**METHODS:** Combining neurolipidomics in patient specimens with functional studies in mouse models, we interrogated apoE isoform-specific functions for sortilin in brain lipid metabolism and AD.

**RESULTS:** Sortilin directs uptake and conversion of polyunsaturated fatty acids into endocannabinoids, lipid-based neurotransmitters that act through nuclear receptors to sustain neuroprotective gene expression in the brain. This sortilin function requires apoE3, but is disrupted by binding of apoE4, impairing endocannabinoid signaling and increasing amyloidogenic processing.

**DISCUSSION:** We uncovered the significance of neuronal apoE receptor sortilin in facilitating neuroprotective actions of brain lipids, and its relevance for AD risk seen with apoE4.

## BACKGROUND

Apolipoprotein (apo) E is the major carrier for lipids in the brain. It is secreted by astrocytes and microglia and delivers essential lipids to neurons that take up apoE-bound cargo via apoE receptors (reviewed in [1]). Apart from its role in lipid homeostasis, apoE also bears significance as the most important genetic risk factor for the sporadic form of Alzheimer’s disease (AD) as carriers of the *APOE4* gene variant are at a significantly higher risk of AD than carriers of the common *APOE3* allele [2]. A large body of work confirmed the potential of apoE4, as compared with apoE3, to promote brain deposition of amyloid-β peptides (Aβ), a major cause of neurodegeneration (reviewed in [3]). Many hypotheses have been advanced how apoE may affect brain Aβ levels, including the delivery of bound Aβ to cellular catabolism via apoE receptors. Still, the mechanism(s) that distinguish apoE3 and apoE4 functions in brain lipid homeostasis and progression of AD remain controversial.

Previously, we identified the lipoprotein receptor sortilin as a major endocytic route for uptake of apoE-containing lipoproteins in neurons *in vitro* and *in vivo*. In gene-targeted mice, loss of sortilin impaired neuronal clearance of murine apoE and was associated with enhanced accumulation of Aβ peptides and senile plaque formation [4]. However, the relevance of the sortilin-dependent uptake of apoE for brain lipid homeostasis and for the risk of AD seen in carriers of the human *APOE4* genotype remained unclear.

Combining MS-based lipidomics in patient specimens with functional studies in humanized mouse models expressing apoE3 or apoE4, we uncovered a unique role for sortilin and apoE3 in facilitating the neuronal metabolism of polyunsaturated fatty acids (PUFA) into endocannabinoids (eCB) that signal an anti-inflammatory gene expression profile in the brain. The ability of sortilin to sustain neuroprotective eCB signaling is disrupted by binding of apoE4, increasing pro-inflammatory markers and aggravating the amyloidogenic burden in the brain.

## METHODS

### Materials and general methods

AD specimens were collected from donors from whom written informed consent for the use of the material for research purposes had been obtained (see supplementary methods). Mouse strains and details on animal experimentation are also given in supplementary methods. Determination of transcript or protein levels in tissue were performed by qRT-PCR and SDS-PAGE, respectively, using standard protocols. Cell culture experiments involving CHO cells are detailed in supplementary methods.

### Lipid analyses

Quantification of levels of total cholesterol, triglycerides, and free fatty acids in plasma was performed using commercial kits (Biovision, Roche, Cayman). Mouse plasma lipoprotein profiles were established by TNO Biosciences (Leiden, The Netherlands) using FPLC. Targeted lipidomics was performed on brain cortex specimens from human subjects or mice or on apoE-containing lipoproteins from human CSF or mouse plasma using LC-MS as detailed in supplementary methods.

## RESULTS

### Sortilin interacts with apoE3 but not apoE4, to reduce brain Aβ42 levels

In mice, inactivation of *Sort1*, the gene encoding sortilin, results in brain accumulation of murine apoE due to impaired clearance of the protein by neurons lacking this apoE receptor [4]. Accumulation of murine apoE coincides with increased levels of Aβ40, suggesting a role for sortilin in neuronal catabolism of apoE-bound Aβ peptides [4]. While these findings implicated sortilin in apoE-dependent control of brain Aβ levels, they failed to address the relevance of this receptor for the increase in Aβ and the risk of AD seen in humans expressing apoE4 as compared with apoE3.

Now, we introduced the *Sort1* defect (KO) into mice carrying a targeted replacement of the murine *Apoe* locus with genes encoding human apoE3 (E3; KO) or apoE4 (E4; KO) [5]. As shown for murine apoE before [4], loss of sortilin also caused accumulation of human apoE3 (Fig. 1A-B) and apoE4 (Fig. 1A, C) in cortex and hippocampus of KO mice compared with wild-types (WT). Sortilin deficiency did not impact transcript levels for apoE3 or apoE4 in the brain (Fig. 1D), nor the levels of apoE3 and apoE4 protein (Fig. S1A) or transcripts (Fig. S1B) in primary astrocytes from (E3;KO) or (E4;KO) mice as compared with WTs. Thus, increased brain levels of human apoE3 and apoE4 in KO mice were likely due to impaired neuronal clearance rather than increased astrocytic production of the apolipoproteins. These findings substantiated sortilin as major clearance receptor for murine and human apoE variants alike.

**Figure 1.**
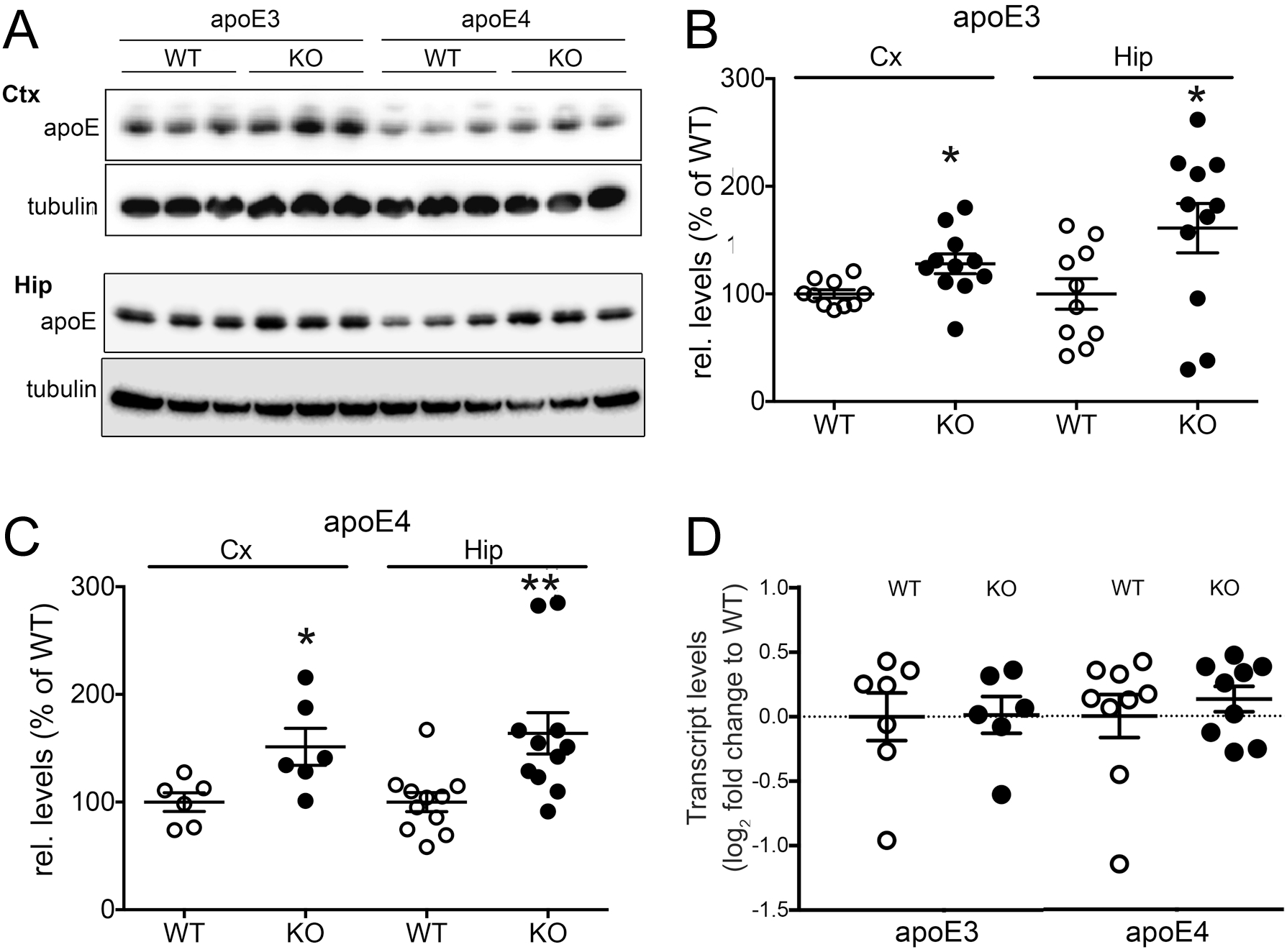
Sortilin deficiency causes brain accumulation of apoE3 and apoE4. (**A-C**) Levels of apoE3 and apoE4 in cortex (Ctx) and hippocampus (Hip) of sortilin WT and KO mice (n=6-11 mice per group) were determined by western blot analysis (A) and densitometric scanning of replicate blots (B and C). Values are mean ± SEM given as percent of WT control (mean set to 100%). Accumulation of apoE3 and apoE4 in KO as compared with WT tissues was determined by Student’s *t* test (*, p<0.05; **, p<0.01). Detection of tubulin served as loading control in A. (**D**) Quantitative RT-PCR analysis of transcript levels for apoE3 and apoE4 in brain extracts of sortilin WT and KO mice at 3 months of age (n=6-9 mice per group). Values are mean ± SEM given as log_2_ fold change compared with levels in the respective WT animals (mean set to 0).

To test the impact of sortilin-mediated clearance of human apoE on brain Aβ levels, we crossed our mouse lines with the PDAPP strain expressing the human amyloid precursor protein APP^V717F^ [6]. Brain accumulation of apoE3 or apoE4 in sortilin-deficient mice did not alter APP levels (Fig. 2A) but coincided with two-fold increased levels of Aβ40 in (E3;KO) and (E4;KO) mice compared with their respective controls (Fig. 2B). This observation supported a role for sortilin in apoE-dependent clearance of Aβ that also extends to human apoE variants. While no difference in accumulation of Aβ40 was evident comparing (E3;KO) and (E4;KO) mice (Fig. 2B), the situation was different for the more pathogenic variant Aβ42. Aβ42 levels were lower in brain cortex of WT mice expressing apoE3 as compared with apoE4, mirroring the higher prevalence of AD in humans with the *APOE4* genotype (Fig. 2C). Loss of sortilin raised Aβ42 levels significantly in mice expressing apoE3 (E3;KO) but did not further increase the already high Aβ42 levels in (E4;KO) animals (Fig. 2C). The increase in Aβ42 in (E3;KO) animals was accompanied by a rise in sAPPβ levels (Fig. 2D), suggesting accelerated proteolysis of APP to sAPPβ and Aβ42 as the cause of increased accumulation of Aβ42 in apoE3 mice lacking sortilin. The same findings were obtained for Aβ40 (Fig. S2B) and Aβ42 (Fig. S2C) in the hippocampi of these mice.

**Figure 2.**
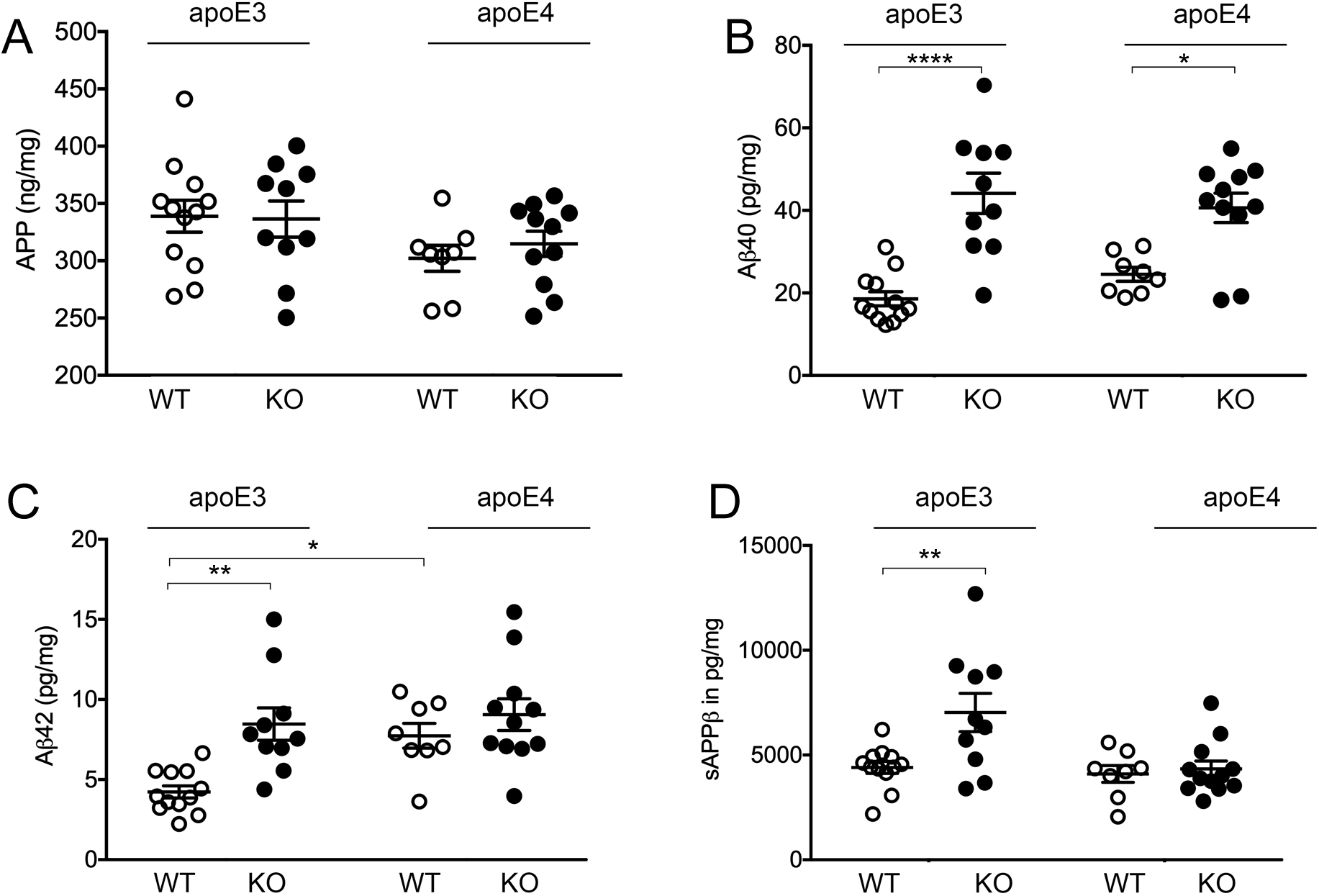
Sortilin and apoE3 control Aβ42 levels in mouse brain cortex. Levels of full-length APP (**A**), soluble Aβ40 (**B**), soluble Aβ42 (**C**), and sAPPβ (**D**) were determined by ELISA in cortical brain extracts of apoE3 or apoE4 targeted replacement mice crossed with the PDAPP strain of mice (4 months of age). In addition, the animals were either wild-type (WT) or homozygous for the *Sort1* null allele (KO). Values are mean ± SEM (n = 8-12 animals per group). Significant differences in levels of APP processing products were tested by Two-way ANOVA, followed by Bonferroni post-hoc analysis. (*, p<0.05; **, p<0.01; ****, p<0.0001).

Taken together, we documented a dual role for sortilin in control of amyloidogenic burden in the brain. It acts as a clearance receptor for Aβ peptides bound to apoE. This pathway regulates brain levels of Aβ40 but does not discriminate between apoE3 or apoE4. However, sortilin also interacts with apoE3 to selectively reduce the amyloidogenic processing of APP to Aβ42. This protective mechanism, that may decrease Aβ42 production by direct or indirect means, is lost with apoE4.

### Interaction of sortilin and apoE3 controls brain levels of PUFAs and endocannabinoids

Next, we queried an interaction of sortilin with apoE in brain lipid homeostasis that may distinguish between apoE3 and apoE4 actions. Our strategy was based on the documented roles for apoE and the lipoprotein receptor sortilin in control of lipid metabolism [4, 7], and on the importance of neuronal lipid homeostasis for risk of AD (reviewed in [8]). To assure that alterations in lipids reflected primary mechanisms rather than secondary consequences of amyloidogenesis, this work was performed in mice lacking the PDAPP transgene.

Initially, we used LC-MS-based lipidomics to determine the levels of various lipid classes in brain cortices of (E3;KO) and (E4;KO) mice. When compared with WTs, we observed a distinct impact of genotypes on brain levels of total fatty acids (FA) and ω-3 polyunsaturated fatty acids (PUFAs), with levels being lower in (E3;KO) as compared with (E3;WT) mice (Fig. 3A). By contrast, levels of these lipid were always low in E4 compared with (E3;WT) animals, irrespective of the presence or absence of sortilin. This interaction of *Sort1* and *APOE* in control of brain lipids was seen for ω-3 but not for ω-6 PUFAs (Fig. 3A), in line with the neuroprotective actions of ω-3 PUFAs in AD [9-11].

**Figure 3.**
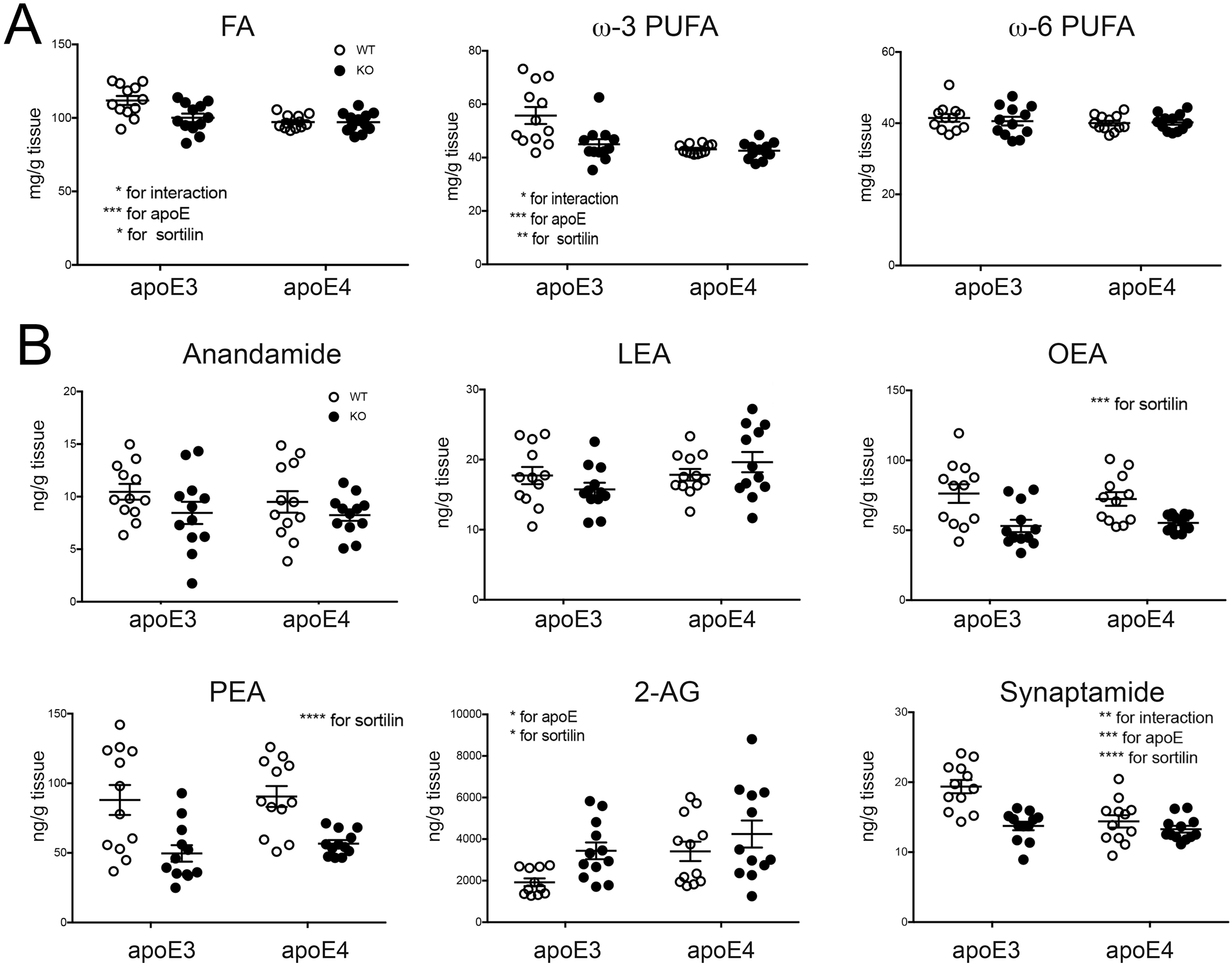
Sortilin and apoE3 interact in control of brain levels of polyunsaturated fatty acids and endocannabinoids. (**A)** Levels of total fatty acids (FA) as well as of ω-3 and ω-6 polyunsaturated fatty acids (PUFA) were determined in brain cortices of apoE3- and apoE4-targeted replacement mice either wild-type (WT) or homozygous for the *Sort1* null allele (KO) (3 months of age, n=12 per genotype). (**B**) Levels of endocannabinoids and endocannabinoid-like lipids were determined in brain cortices of mice of the indicated *APOE* and *Sort1* genotypes (3 months of age, n=11-12 per genotype). Data are the mean ± SEM. The significance of data was determined by Two-way ANOVA, followed by Bonferroni post-hoc analysis (*, p<0.05; **, p<0.01; ***, p<0.001; ****, p<0.0001). 2-AG, 2-arachidonoylglycerol; LEA, linoleoyl-ethanolamide; OEA, oleyl-ethanolamide; PEA, palmitoyl-ethanolamide.

To delineate the relevance of sortilin and apoE3 for brain PUFA metabolism, we determined levels of bioactive PUFA derivatives in brains of our four mouse strains (Fig. 3B). We focused on endocannabinoids (eCBs), lipid-based neurotransmitters produced from PUFAs in neurons. Most eCBs act anti-inflammatory and neuroprotective, and alterations in eCB levels have been associated with AD [12]. Of the six eCBs and related lipids tested (Fig. 3B), levels of anandamide and linoleoyl-ethanolamide (LEA) were not impacted by genotypes. By contrast, levels of oleyl-ethanolamide (OEA) and palmitoyl-ethanolamide (PEA) were reduced in animals lacking sortilin, but irrespective of apoE genotype. Finally, levels of 2-arachidonoylglycerol (2-AG) and the eCB-like lipid synaptamide showed interaction between *APOE* and *Sort1* as observed for brain Aβ42 levels before. In detail, physiological 2-AG levels were low in (E3;WT) mice but increased in (E3;KO) animals. By contrast, levels in E4 mice were always high compared with (E3;WT), irrespective of *Sort1* genotype. The converse pattern was seen for synaptamide with physiological levels being high in (E3;WT) but lower in all other genotype combinations. The dependency of eCB levels on *APOE* genotype was substantiated in brain specimens of AD patients homozygous for *APOE3* or *APOE4* (Fig. 4). As in mice, levels of synaptamide were lower (p<0.05) while levels of 2-AG tended to be higher (p=0.057) in *APOE4* as compared with *APOE3* carriers. Levels of anandamide were also decreased in the *APOE4* genotype (p<0.05), whereas OEA, LEA, and PEA levels were not impacted.

**Figure 4.**
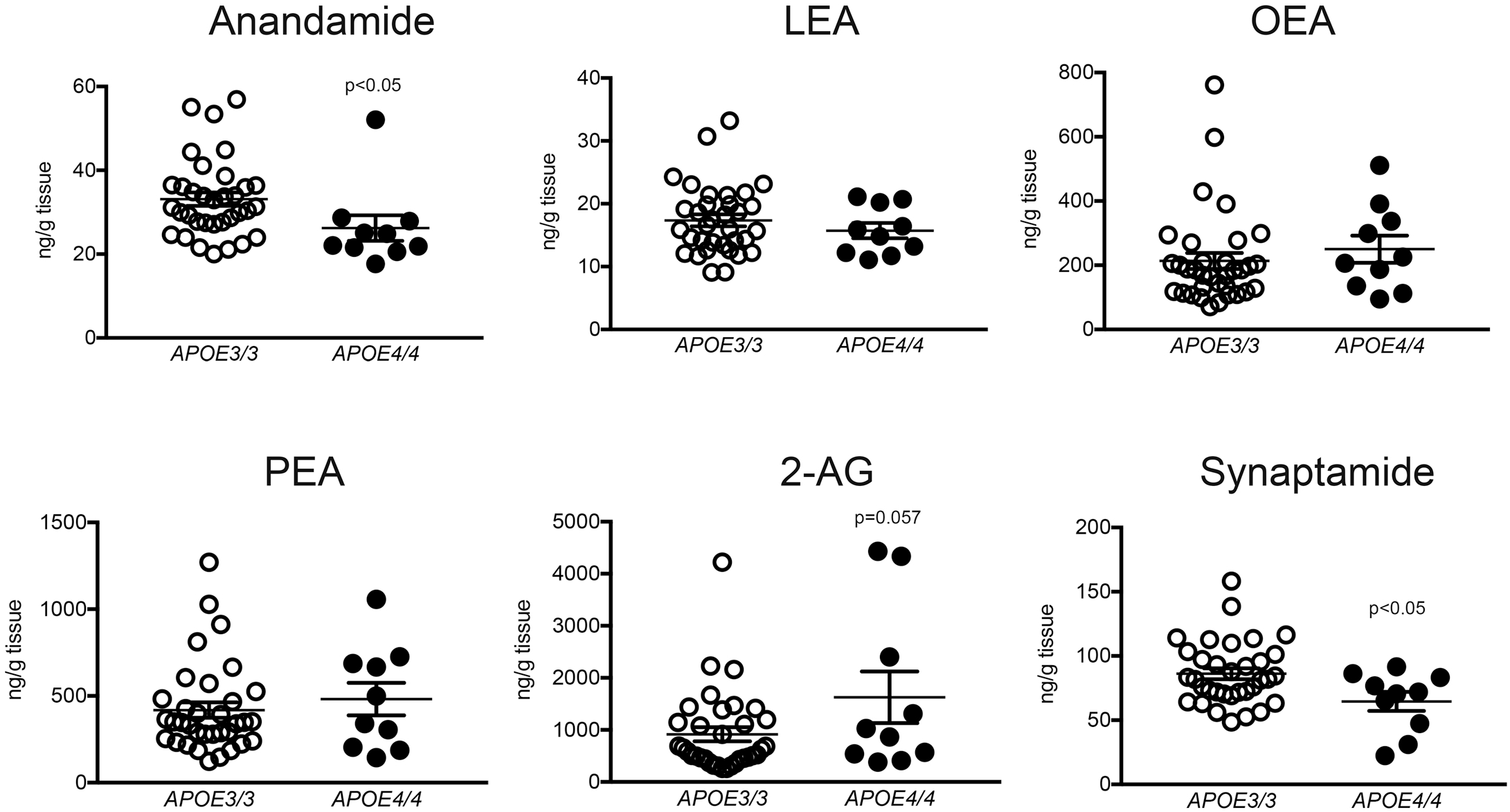
ApoE-dependent changes in eCB levels in the human brain. Levels of the indicated eCBs and eCB-like lipids were determined in prefrontal cortex specimens of AD patients homozygous for *APOE3* or *APOE4* (n=10 for *APOE4*, n=34 for *APOE3*.) Values are mean ± SEM. The significance of data was determined by unpaired Student *t* test. 2-AG, 2-arachidonoylglycerol; LEA, linoleoyl-ethanolamide; OEA, oleyl-ethanolamide; PEA, palmitoyl-ethanolamide.

### *Sort1* and *APOE3* safeguard an anti-inflammatory gene expression profile signaled by PPARs

Endocannabinoids exert their actions by signaling via G-protein coupled cannabinoid receptors CB1 and CB2 or by acting as ligands for peroxisome proliferator-activated receptors (PPARs). Because synaptamide does not engage CB1/2 [13], we explored the relevance of sortilin and apoE3 interaction in eCB metabolism by focusing on PPARs. Using a microarray-based strategy to assess transcript levels of 84 PPAR target genes, we identified 12 brain transcripts that showed dependence on *Sort1* and *APOE*, being either higher or lower in (E3;WT) as compared with the other three genotypes (Fig. 5A). Affected transcripts included PPARγ (*Pparg)* and the retinoic X receptor (*Rxrg*), which forms heterodimers with PPARs; but also factors in lipid homeostasis, such as fatty acid transporter Slc27a4 and very long-chain acyl-CoA synthetase Slc27a5. Using qRT-PCR for PPAR targets *Pparg, Mmp9*, and *Klf10*, we substantiated an effect of sortilin on eCB-dependent gene transcription in E3 but not in E4 mice (Fig. 5B). PPARs are lipid sensors that suppress inflammatory responses by inducing an anti-inflammatory gene expression profile, an activity reducing neurodegeneration [14, 15]. In line with interaction of *Sort1* and *APOE3* in promoting PPAR activities, loss of sortilin in apoE3 mice or the presence of apoE4 (irrespective of *Sort1* genotype) resulted in alterations in the mouse brains consistent with a pro-inflammatory state. These changes included decreased transcription of *Vegf*, but elevated transcript levels of *Tnfα* and *Gfap* in (E3;KO), (E4;WT), and (E4;KO) mice as compared with (E3;WT) (Fig. 5C). A potential pro-inflammatory state in the three mouse strains as compared with (E3;WT) was supported by increased immunosignals for GFAP in the brain (Fig. 5D-E).

**Figure 5.**
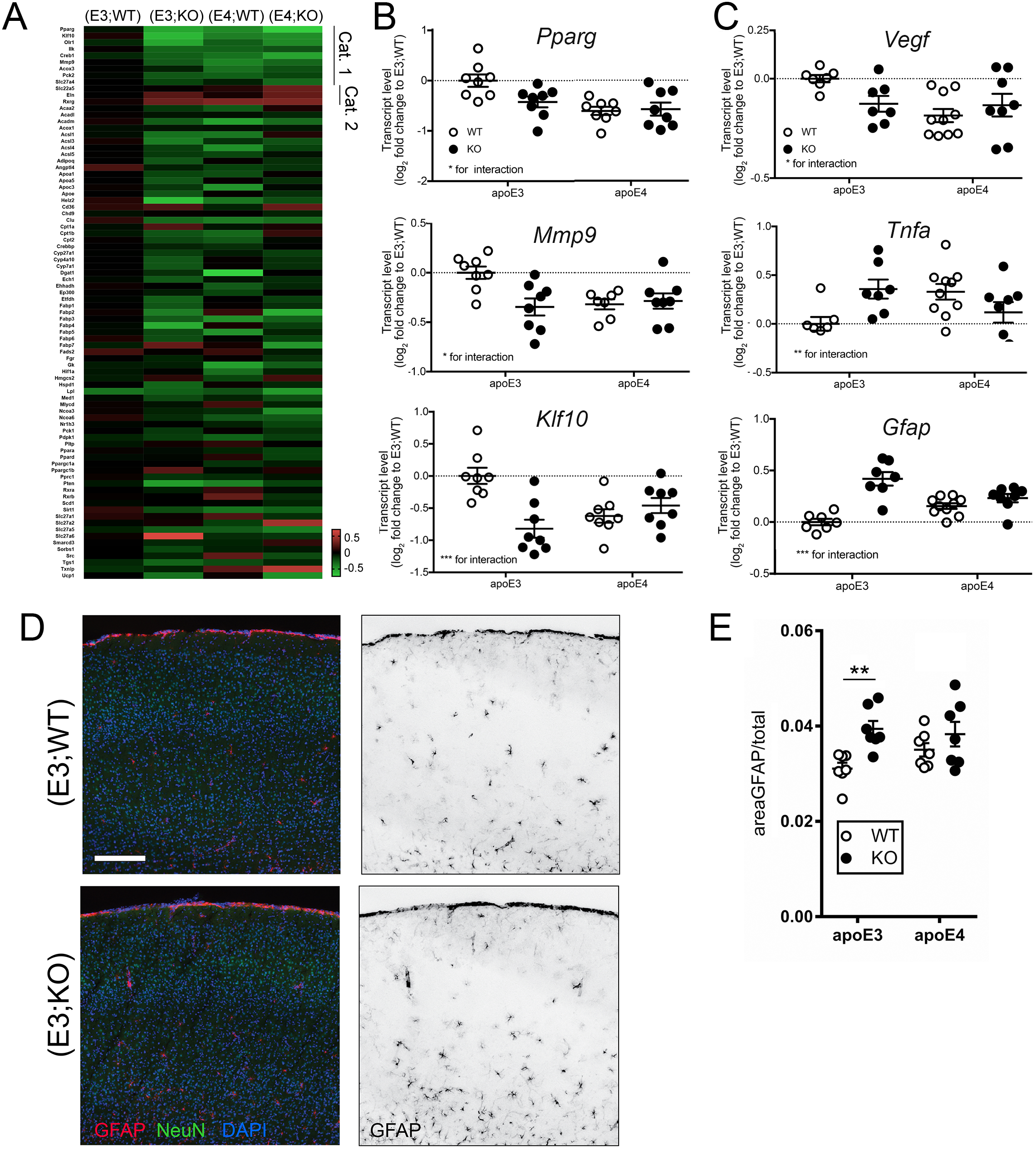
Interaction of *Sort1* and *APOE3* controls PPAR-dependent gene expression. (**A**) Heatmap of expression levels of target genes of peroxisome proliferator-activated receptors (PPAR) in the brain of apoE3 or apoE4 targeted replacement mice either wild-type (WT) or homozygous for the *Sort1* null allele (KO) (Mouse PPAR Targets RT^2^ Profiler PCR Array, Qiagen). Transcripts up (Cat. 1) or down (Cat. 2) in (E3;WT) as compared with the other three genotype groups are highlighted (n=3-4 mice per group). (**B**) Quantitative RT-PCR analysis of transcript levels of the indicated PPAR target genes in brain cortices of apoE3 and apoE4 mice either WT or KO for *Sort1*. Log_2_ fold changes as compared to (E3;WT) mice set to value 0 are given (n=7-8 mice per group; Two-way ANOVA). (**C**) Quantitative RT-PCR analysis of vascular endothelial growth factor (*Vegf*), tumor necrosis factor α (*Tnfa*), and glial fibrillary acidic protein (*Gfap*) in brain cortices of mice of the indicated *Sort1* and *APOE* genotypes. Log_2_ fold changes as compared to (E3;WT) mice set to value 0 are given (n=7-10 mice per group; Two-way ANOVA). Mice were 70-75 weeks of age. (**D**) Immunodetection of GFAP (red) on exemplary cortical brain sections of (E3;WT) and (E3;KO) mice. Sections were also stained for neuronal marker NeuN (green) and DAPI (blue). Both merged color images (left panels) and single GFAP channels in grey scale (right panels) are given. Scale bar: 250 µm. (**E**) Quantification of the GFAP-immunoreactive area in the cortex of apoE3 and apoE4 mice either WT or KO for *Sort1* is shown (n=7 mice per group; Two-way ANOVA with Tukey’s multiple comparisons test). Mice were 70-75 weeks of age. *, *p*<0.05; **, *p*<0.01; ***, p<0.001.

Our data indicated interaction of *Sort1* and *APOE3* in safeguarding a neuroprotective metabolism and action of eCBs, potentially protecting the brain from inflammatory insults and decreasing amyloidogenic stimuli. This neuroprotective eCB action is lost in apoE3 mice that lack sortilin, raising brain Aβ42 levels. By contrast, this neuroprotective action of sortilin in brain lipid metabolism is not supported by apoE4, as PPAR activities are decreased and Aβ42 levels increased in E4 mice irrespective of *Sort1* genotype.

### Sortilin and apoE3 do not impact systemic metabolism of PUFAs

Anandamide and 2-AG are produced from arachidonic acid (ARA), while synaptamide is derived from docosahexaenoic acid (DHA). Mainly, ARA and DHA are supplied to the brain from the blood stream as FA or esterified to phospholipids. In the brain, ARA and DHA are re-esterified to membrane phospholipids or converted into bioactive metabolites, such as eCBs (reviewed in [16]). To query the impact of sortilin and apoE on PUFA transport into the brain, we quantified lipid levels in brain tissue, and in plasma and brain lipoproteins. Levels of DHA, but not of ARA, were reduced in the brains of (E3;KO), (E4;WT), and (E4;KO) compared with (E3;WT) mice (Fig. S3A). A similar trend in reduction of DHA (p=0.1) but not ARA levels was seen in brain tissues from AD patients with *APOE4/4* as compared with *APOE3/3* (Fig. S3B). Altered brain levels of DHA were not due to differential association of lipids with lipoproteins containing apoE3 or apoE4 as plasma lipoprotein profiles were indistinguishable comparing all four mouse genotypes (Fig. S4A), as were the levels of DHA and ARA in apoE-containing lipoproteins isolated from plasma of these mice (Fig. S4B). Also, levels of DHA and ARA were comparable in apoE3- and apoE4-containing brain lipoproteins isolated from human cerebrospinal fluid (Fig. S4C-F). Finally, transcript levels of enzymes in neuronal metabolism of 2-AG and synaptamide were not affected by *Sort1* or *APOE* genotype as shown by qPCR on mouse brain extracts (Fig. S5).

### ApoE4 disrupts cell surface recycling of sortilin

So far, our studies failed to identify changes in biosynthesis or extracellular transport that may explain why alterations in DHA and eCB metabolism in E4 mice resembled defects in apoE3 animals lacking sortilin. Thus, we focused on the hypothesis that apoE4 may disrupt the activity of sortilin, rendering E4 brains essentially sortilin-depleted. This hypothesis was based on the propensity of apoE4 to impair intracellular sorting and thereby activity of several cell surface receptors [17-19].

To query whether trafficking of sortilin was differentially impacted by apoE variants, we tested its co-localization with apoE3 or apoE4 in CHO cells using proximity ligation assays (PLA). Co-localization of sortilin with apoE3 was seen in a scattered vesicular pattern throughout the cytoplasm, whereas co-localization with apoE4 showed a distinct pattern close the cell membrane (Fig. 6A). To substantiate this differential impact of apoE4 on sortilin sorting, we established the trafficking path of the receptor in CHO cells by labeling sortilin molecules on the cell surface with antibodies and following their subsequent intracellular route using immunocytochemistry (Fig. 6B). Within 15 minutes, labeled receptors internalized from the cell surface (Fig. 6B, panel surface labeling) into intracellular compartments (Fig. 6B, panel internalization). To interrogate recycling of these internalized receptors, we subsequently treated the cells with dynasore, an inhibitor of endocytosis to block continuous re-entry of recycled receptors into the cells. Application of dynasore resulted in the accumulation of labeled sortilin molecules at the plasma membrane, confirming cell surface recycling of internalized (antibody-labeled) receptors (Fig. 6B, panel recycling). This recycling path of sortilin was not impacted by the presence of apoE3 in the culture medium (Fig. 6C). However, in the presence of apoE4, internalized receptors failed to re-appear on the cell surface and remained largely intracellular (Fig. 6C). To more accurately quantify the extent of sortilin recycling in the presence of apoE3 versus apoE4, we biotinylated sortilin molecules on the surface of CHO cells and followed their internalization and recycling fate as schematized in Fig. 6D. Recycled receptors accumulating at the cell surface were stripped of their biotin tag by glutathione treatment to only retain a biotin label in the intracellular (non-recycling) receptor pool. The extent of receptor recycling was determined by subtracting the amount of biotinylated receptors purified on streptavidin beads after 60 minutes of recycling from the total amount of biotinylated receptors internalized at the beginning of the experiment. In these studies, the amount of receptors recycling back to the cell surface was reduced by 50% in apoE4-treated as compared to apoE3-treated cells (Fig. 6E and F).

**Figure 6.**
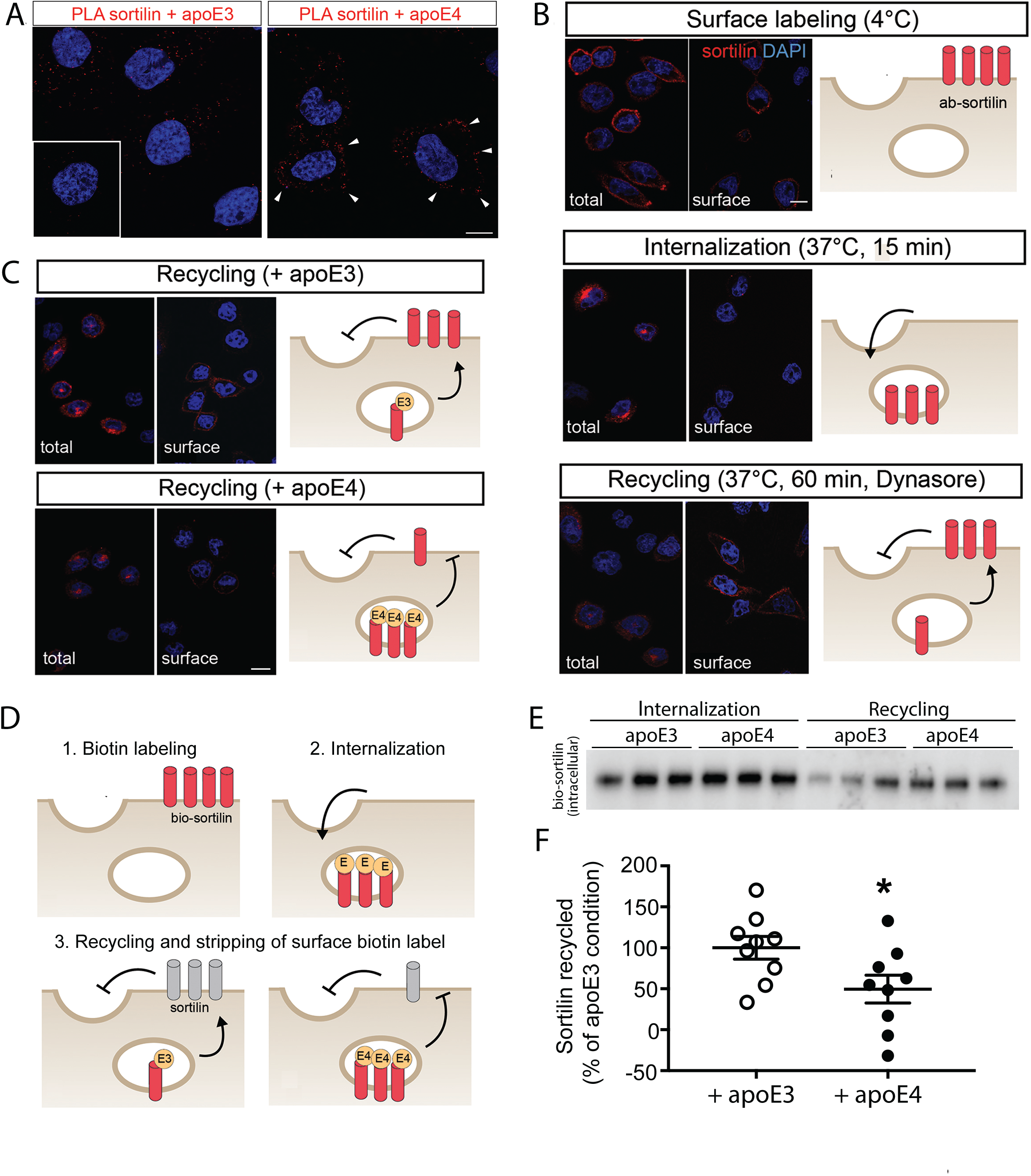
ApoE4 disrupts intracellular trafficking of sortilin. **(A)** CHO cells stably expressing murine sortilin (CHO-S) were treated for 24 hours at 37°C with 5 µg/ml of myc-tagged apoE3 or apoE4 produced in HEK293 cells [4]. Co-localization of sortilin and apoE was tested by proximity ligation assay (PLA) using primary antisera directed against sortilin and myc, respectively. PLA signals (red) for colocalization showed a dispersed vesicular pattern with apoE3 but a juxtamembrane pattern with apoE4 (white arrowheads). The inset documents the absence of PLA signals in cells not treated with apoE. Cell nuclei were counterstained with DAPI (blue). Scale bar: 10 µm. (**B**) Immunodetection of sortilin (red) in CHO-S cells under the indicated treatment conditions. Cells were either permeabilized (total) or non-permeabilized (surface) to distinguish between total and cell surface receptor pools. Cells were counterstained with DAPI (blue). Scale bar: 10 µm. Schematics visualize the proposed trafficking fate of sortilin in the respective images. *Panel Surface labeling*: Receptor molecules at the cell surface were decorated with primary antibody (ab-sortilin, red symbols) in CHO-S cells kept at 4°C to disrupt membrane trafficking. Immunosignals for sortilin in non-permeabilized cells (surface) confirm plasma membrane localization of the labeled receptors. *Panel Internalization*: Following shift to 37°C for 15 minutes, labeled receptors move to an intracellular localization as evidenced by immunosignals in the permeabilized (total) but not in the non-permeabilized (surface) condition. *Panel Recycling*: To confirm re-appearance of internalized receptors at the cell surface, cells were subsequently treated with dynasore for 60 min at 37°C. When endocytosis was blocked with dynasore to prevent re-uptake of surface localized receptors, labeled sortilin molecules accumulated at the cell surface as evidenced by immunosignals in non-permeabilized cells (surface), substantiating a recycling path for the receptor. (**C**) Experiment as in (B) but cells were incubated in apoE3- or apoE4-conditioned medium. Antibody-tagged sortilin molecules recycled to cell surface in the presence of apoE3 as documented by immunosignals in non-permeabilized cells (upper panels). No receptor recycling was observed in the presence of apoE4 as evidenced by the absence of immunosignals in the non-permeabilized condition (lower panels). (**D**) Experimental strategy to quantify sortilin recycling in the presence of apoE3 or apoE4. Sortilin molecules at the surface of CHO-S cells kept at 4°C were biotinylated (panel Labeling; bio-sortilin, red receptor symbols). Following shift to 37°C for 15 minutes in the presence of 5 µg/ml of apoE3 or apoE4 (yellow symbols), receptors remaining at the cell surface were stripped of their biotin label using glutathione (GSH) treatment (panel 2, Internalization). Finally, cells were incubated for 60 minutes at 37°C in the presence of dynasore and then with GSH, stripping all recycled receptors accumulating at the surface of biotin label (panel 3, Recycling; gray receptor symbols). (**E**) Western blot analysis of biotinylated sortilin molecules affinity-purified on streptavidin beads from CHO-S cells after internalization and recycling phases as described in (D, steps 2 and 3). (**F**) The amount of biotinylated (bio-)sortilin was determined by densitometric scanning of replicate western blots as exemplified in (E). The % of recycled receptors was determined by subtracting the amount of biotinylated (intracellular) receptors at the recycling phase (step 3) from that at internalization (step 2). Mean values for apoE3 were set to 100%. Significantly less receptors recycled in the presence of apoE4 than apoE3 (49.56 ± 16.92% vs. 100 ± 13.84%). (data are given as mean ± SEM; n= 9 biological replicates from three independent experiments, Student’s *t* test; *, <0.05)

## DISCUSSION

We propose a novel concept for the neuroprotective metabolism of PUFA in the brain. It involves interaction of sortilin with apoE3 to support neuronal uptake and action of DHA and eCBs, providing anti-inflammatory gene expression und suppressing noxious insults that increase Aβ42 production. Loss of sortilin in apoE3 mice compromises this neuroprotective PUFA uptake pathway, increasing pro-inflammatory gene expression and amyloidogenic processing. Similar detrimental effects are seen in wild-type mice in the presence of apoE4 as it disrupts receptor trafficking.

In mice, loss of sortilin reduces neuronal uptake of apoE and causes alterations in brain lipid homeostasis also seen in apoE KO animals (i.e., accumulation of sulfatides) [4]. These data suggested sortilin as a key player in apoE-dependent lipid metabolism in the brain. Our new results substantiate this role by identifying the interaction of sortilin with apoE3 in control of neuronal PUFA metabolism and signaling. Likely, this neuroprotective lipid pathway explains the anti-amyloidogenic interaction of apoE3 and sortilin (Figs. 2 and S2) as the same interaction between *Sort1* and *APOE* genotypes, as for Aβ42, is also seen for brain levels of DHA and several eCBs (Fig. 3). Importantly, the same effect of *APOE4* on DHA and eCBs levels is seen in AD patients, substantiating the clinical relevance of our findings from mouse models (Fig. 4).

DHA and its bioactive metabolites are regulators of brain health, linking inflammation with neurodegeneration. DHA is the most abundant brain ω-3 PUFA. It regulates numerous cellular processes, especially the resolution of brain inflammation [16, 20]. Low levels of DHA in plasma or brain correlate with human AD [21-23] and dietary DHA intake lowers Aβ levels in mice [9-11]. A similar role in control of inflammation in the ageing brain has been suggested for anandamide and 2-AG. Anandamide levels decrease in the brain of patients [24, 25] and mice with AD [26]. Conversely, levels of 2-AG increase in AD patients [27] and mouse models [28], and increased 2-AG signaling exacerbates synaptic failure [29]. Although little is known about synaptamide, it is considered a bioactive mediator of DHA as it recapitulates the protective actions seen with DHA [30]. We document a pathological increase in brain levels of 2-AG and a concomitant decrease in DHA, anandamide, and synaptamide in humans and mice with apoE4, and in apoE3 mice lacking sortilin, identifying the relevance of *Sort1* and *APOE* genotypes for brain PUFA homeostasis.

We focused on the relevance of sortilin and apoE3 interaction for DHA and eCB signaling by investigating PPARs. Transcript levels of PPAR-γ increase in AD patients to counteract neuroinflammatory insults [31]. Non-steroidal anti-inflammatory drugs ameliorate AD-related processes due to their ability to stimulate PPAR and inhibit inflammatory responses [32, 33]. In line with a function for sortilin and apoE3 in PPAR actions, we observed a concordant dysregulation of transcriptional targets in E4 mice and in E3 animals lacking sortilin (Fig. 5A-B). The relevance of *Sort1* and *APOE3* to counteract brain inflammation was supported by an increase in pro-inflammatory markers GFAP and TNFα (Fig. 5C-E). GFAP indicates brain inflammation in rodent AD models [34], and increases in GFAP levels in AD rats are reverted by eCB application [35]. Increased levels of TNFα are also linked to AD pathology [36, 37]. Another sign of a potentially pro-inflammatory state in E4 mice and in (E3;KO) animals is down-regulation of VEGF (Fig. 5C). VEGF suppresses inflammatory brain response [38] and reduces neurodegeneration in mouse models of AD [39, 40].

Sortilin facilitates cellular uptake of apoE3 and apoE4 equally well, both *in vitro* [4] and *in vivo* (Fig. 1A-C). Also, levels of PUFA bound to apoE3- or apoE4-containing lipoprotein in plasma or brain are comparable (Fig. S4). Thus, apoE variants likely do not discriminate the types of PUFA taken up into neurons via sortilin. Rather it is the detrimental effect of apoE4 on trafficking of sortilin that impairs this neuronal uptake route for lipids (Fig. 6). Following endocytic uptake, apoE3 recycles to the cell surface for re-secretion. This route is not taken by apoE4 that accumulates in the endocytic pathway [17, 18, 41]. Impaired recycling of apoE4 disrupts cell surface re-exposure of apoE binding proteins, as shown for APOER2 [19] or the insulin receptor [42], and reversal of impaired recycling has been suggested as therapeutic approach for AD risk imposed by apoE4 [43]. We show a similar deleterious impact of apoE4 on sortilin trafficking (Fig. 6), likely explaining the loss of neuroprotective actions of this receptor in eCB metabolism and Aβ42 accumulation in the apoE4 brain.

In conclusion, we identify a novel pathway linking neuronal apoE handling by sortilin with the well-known neuroprotective actions of PUFA in the brain, and we provide a possible molecular explanation for the risk of AD seen with apoE4.

## Acknowledgement

We are indebted to T. Pasternack, K. Kampf, C. Kruse, M. Kahlow, and S. Ehret for expert technical assistance. Studies were funded in part by the ERC (BeyOND No. 335692), the Helmholtz Association (AMPro), the Alzheimer Forschung Initiative (#18003), and the Novo Nordisk Foundation (NNF18OC0033928) to TEW.

## REFRENCES

1. Holtzman DM, Herz J, Bu G. Apolipoprotein e and apolipoprotein e receptors: normal biology and roles in Alzheimer disease. Cold Spring Harb Perspect Med. 2012;2:a006312.

2. Corder EH, Saunders AM, Strittmatter WJ, Schmechel DE, Gaskell PC, Small GW, et al. Gene dose of apolipoprotein E type 4 allele and the risk of Alzheimer’s disease in late onset families. Science. 1993;261:921–3.

3. Liu CC, Liu CC, Kanekiyo T, Xu H, Bu G. Apolipoprotein E and Alzheimer disease: risk, mechanisms and therapy. Nat Rev Neurol. 2013;9:106–18.

4. Carlo AS, Gustafsen C, Mastrobuoni G, Nielsen MS, Burgert T, Hartl D, et al. The pro-neurotrophin receptor sortilin is a major neuronal apolipoprotein E receptor for catabolism of amyloid-beta peptide in the brain. The Journal of neuroscience: the official journal of the Society for Neuroscience. 2013;33:358–70.

5. Knouff C, Hinsdale ME, Mezdour H, Altenburg MK, Watanabe M, Quarfordt SH, et al. Apo E structure determines VLDL clearance and atherosclerosis risk in mice. J Clin Invest. 1999;103:1579–86.

6. Games D, Adams D, Alessandrini R, Barbour R, Berthelette P, Blackwell C, et al. Alzheimer-type neuropathology in transgenic mice overexpressing V717F beta-amyloid precursor protein. Nature. 1995;373:523–7.

7. Huang Y, Mahley RW. Apolipoprotein E: structure and function in lipid metabolism, neurobiology, and Alzheimer’s diseases. Neurobiol Dis. 2014;72 Pt A:3–12.

8. Di Paolo G, Kim TW. Linking lipids to Alzheimer’s disease: cholesterol and beyond. Nat Rev Neurosci. 2011;12:284–96.

9. Calon F, Lim GP, Yang F, Morihara T, Teter B, Ubeda O, et al. Docosahexaenoic acid protects from dendritic pathology in an Alzheimer’s disease mouse model. Neuron. 2004;43:633–45.

10. Lim GP, Calon F, Morihara T, Yang F, Teter B, Ubeda O, et al. A diet enriched with the omega-3 fatty acid docosahexaenoic acid reduces amyloid burden in an aged Alzheimer mouse model. J Neurosci. 2005;25:3032–40.

11. Green KN, Martinez-Coria H, Khashwji H, Hall EB, Yurko-Mauro KA, Ellis L, et al. Dietary docosahexaenoic acid and docosapentaenoic acid ameliorate amyloid-beta and tau pathology via a mechanism involving presenilin 1 levels. J Neurosci. 2007;27:4385–95.

12. Bedse G, Romano A, Lavecchia AM, Cassano T, Gaetani S. The role of endocannabinoid signaling in the molecular mechanisms of neurodegeneration in Alzheimer’s disease. J Alzheimers Dis. 2015;43:1115–36.

13. Sheskin T, Hanus L, Slager J, Vogel Z, Mechoulam R. Structural requirements for binding of anandamide-type compounds to the brain cannabinoid receptor. J Med Chem. 1997;40:659–67.

14. O’Sullivan SE. An update on PPAR activation by cannabinoids. Br J Pharmacol. 2016;173:1899–910.

15. Heneka MT, Reyes-Irisarri E, Hull M, Kummer MP. Impact and Therapeutic Potential of PPARs in Alzheimer’s Disease. Curr Neuropharmacol. 2011;9:643–50.

16. Bazinet RP, Laye S. Polyunsaturated fatty acids and their metabolites in brain function and disease. Nat Rev Neurosci. 2014;15:771–85.

17. Heeren J, Beisiegel U, Grewal T. Apolipoprotein E recycling: implications for dyslipidemia and atherosclerosis. Arterioscler Thromb Vasc Biol. 2006;26:442–8.

18. Heeren J, Grewal T, Laatsch A, Becker N, Rinninger F, Rye KA, et al. Impaired recycling of apolipoprotein E4 is associated with intracellular cholesterol accumulation. J Biol Chem. 2004;279:55483–92.

19. Chen Y, Durakoglugil MS, Xian X, Herz J. ApoE4 reduces glutamate receptor function and synaptic plasticity by selectively impairing ApoE receptor recycling. Proc Natl Acad Sci U S A. 2010;107:12011–6.

20. Lukiw WJ, Cui JG, Marcheselli VL, Bodker M, Botkjaer A, Gotlinger K, et al. A role for docosahexaenoic acid-derived neuroprotectin D1 in neural cell survival and Alzheimer disease. J Clin Invest. 2005;115:2774–83.

21. Schaefer EJ, Bongard V, Beiser AS, Lamon-Fava S, Robins SJ, Au R, et al. Plasma phosphatidylcholine docosahexaenoic acid content and risk of dementia and Alzheimer disease: the Framingham Heart Study. Arch Neurol. 2006;63:1545–50.

22. Soderberg M, Edlund C, Kristensson K, Dallner G. Fatty acid composition of brain phospholipids in aging and in Alzheimer’s disease. Lipids. 1991;26:421–5.

23. Yuki D, Sugiura Y, Zaima N, Akatsu H, Takei S, Yao I, et al. DHA-PC and PSD-95 decrease after loss of synaptophysin and before neuronal loss in patients with Alzheimer’s disease. Sci Rep. 2014;4:7130.

24. Jung KM, Astarita G, Yasar S, Vasilevko V, Cribbs DH, Head E, et al. An amyloid beta42-dependent deficit in anandamide mobilization is associated with cognitive dysfunction in Alzheimer’s disease. Neurobiol Aging. 2012;33:1522–32.

25. Koppel J, Bradshaw H, Goldberg TE, Khalili H, Marambaud P, Walker MJ, et al. Endocannabinoids in Alzheimer’s disease and their impact on normative cognitive performance: a case-control and cohort study. Lipids Health Dis. 2009;8:2.

26. Maroof N, Ravipati S, Pardon MC, Barrett DA, Kendall DA. Reductions in endocannabinoid levels and enhanced coupling of cannabinoid receptors in the striatum are accompanied by cognitive impairments in the AbetaPPswe/PS1DeltaE9 mouse model of Alzheimer’s disease. J Alzheimers Dis. 2014;42:227–45.

27. Altamura C, Ventriglia M, Martini MG, Montesano D, Errante Y, Piscitelli F, et al. Elevation of Plasma 2-Arachidonoylglycerol Levels in Alzheimer’s Disease Patients as a Potential Protective Mechanism against Neurodegenerative Decline. J Alzheimers Dis. 2015;46:497–506.

28. Piro JR, Benjamin DI, Duerr JM, Pi Y, Gonzales C, Wood KM, et al. A dysregulated endocannabinoid-eicosanoid network supports pathogenesis in a mouse model of Alzheimer’s disease. Cell Rep. 2012;1:617–23.

29. Mulder J, Zilberter M, Pasquare SJ, Alpar A, Schulte G, Ferreira SG, et al. Molecular reorganization of endocannabinoid signalling in Alzheimer’s disease. Brain. 2011;134:1041–60.

30. Rashid MA, Katakura M, Kharebava G, Kevala K, Kim HY. N-Docosahexaenoylethanolamine is a potent neurogenic factor for neural stem cell differentiation. J Neurochem. 2013;125:869–84.

31. de la Monte SM, Wands JR. Molecular indices of oxidative stress and mitochondrial dysfunction occur early and often progress with severity of Alzheimer’s disease. J Alzheimers Dis. 2006;9:167–81.

32. Heneka MT, Sastre M, Dumitrescu-Ozimek L, Hanke A, Dewachter I, Kuiperi C, et al. Acute treatment with the PPARgamma agonist pioglitazone and ibuprofen reduces glial inflammation and Abeta1-42 levels in APPV717I transgenic mice. Brain. 2005;128:1442–53.

33. Pedersen WA, McMillan PJ, Kulstad JJ, Leverenz JB, Craft S, Haynatzki GR. Rosiglitazone attenuates learning and memory deficits in Tg2576 Alzheimer mice. Exp Neurol. 2006;199:265–73.

34. Liu CC, Zhao N, Yamaguchi Y, Cirrito JR, Kanekiyo T, Holtzman DM, et al. Neuronal heparan sulfates promote amyloid pathology by modulating brain amyloid-beta clearance and aggregation in Alzheimer’s disease. Sci Transl Med. 2016;8:332ra44.

35. Scuderi C, Stecca C, Valenza M, Ratano P, Bronzuoli MR, Bartoli S, et al. Palmitoylethanolamide controls reactive gliosis and exerts neuroprotective functions in a rat model of Alzheimer’s disease. Cell Death Dis. 2014;5:e1419.

36. Alvarez A, Cacabelos R, Sanpedro C, Garcia-Fantini M, Aleixandre M. Serum TNF-alpha levels are increased and correlate negatively with free IGF-I in Alzheimer disease. Neurobiol Aging. 2007;28:533–6.

37. Zhao M, Cribbs DH, Anderson AJ, Cummings BJ, Su JH, Wasserman AJ, et al. The induction of the TNFalpha death domain signaling pathway in Alzheimer’s disease brain. Neurochem Res. 2003;28:307–18.

38. Xu Z, Han K, Chen J, Wang C, Dong Y, Yu M, et al. Vascular endothelial growth factor is neuroprotective against ischemic brain injury by inhibiting scavenger receptor A expression on microglia. Journal of Neurochemistry. 2017;142:700–9.

39. Herran E, Perez-Gonzalez R, Igartua M, Pedraz JL, Carro E, Hernandez RM. Enhanced Hippocampal Neurogenesis in APP/Ps1 Mouse Model of Alzheimer’s Disease After Implantation of VEGF-loaded PLGA Nanospheres. Curr Alzheimer Res. 2015;12:932–40.

40. Religa P, Cao R, Religa D, Xue Y, Bogdanovic N, Westaway D, et al. VEGF significantly restores impaired memory behavior in Alzheimer’s mice by improvement of vascular survival. Scientific Reports. 2013;3:2053.

41. Heeren J, Grewal T, Laatsch A, Rottke D, Rinninger F, Enrich C, et al. Recycling of apoprotein E is associated with cholesterol efflux and high density lipoprotein internalization. J Biol Chem. 2003;278:14370–8.

42. Zhao N, Liu CC, Van Ingelgom AJ, Martens YA, Linares C, Knight JA, et al. Apolipoprotein E4 Impairs Neuronal Insulin Signaling by Trapping Insulin Receptor in the Endosomes. Neuron. 2017;96:115–29 e5.

43. Xian X, Pohlkamp T, Durakoglugil MS, Wong CH, Beck JK, Lane-Donovan C, et al. Reversal of ApoE4-induced recycling block as a novel prevention approach for Alzheimer’s disease. Elife. 2018;7.

