## supplementary methods and figures for "Apolipoprotein E4 disrupts the neuroprotective action of sortilin in neuronal lipid metabolism and endocannabinoid signaling"

### SUPPLEMENTAL METHODS

#### Human specimens

Brain autopsy specimens from the frontal cortex of AD patients and control subjects were obtained from the Netherlands Brain Bank (Netherlands Institute for Neuroscience, Amsterdam) and the MRC London Brain Bank for Neurodegenerative Diseases (Institute of Psychiatry, King's College London). The ethnicity of samples was white. All material was collected from donors for or from whom a written informed consent for a brain autopsy and the use of the material and clinical information for research purposes had been obtained by the Netherlands Brain Bank or the MRC London Brain Bank. Detailed personal information, including age, gender, neuropathological stage, and *APOE* genotype, of the individuals were provided by the brain banks. Cerebrospinal fluid (CSF) samples from AD patients and controls were obtained from a local biomaterial bank. The study was approved by an institutional review board (Ethikausschuss 2 Charité; BIH CRG 2a, EA2/118/15). Written informed consent was obtained from all individuals prior to participating in this study.

#### Animal experimentation

The generation of PDAPP [1], *Sort1*$^{-/-}$ [2], and *Apoe* targeted replacement [3] strains of mice has been described. The animals were kept on a normal chow (4.5% crude fat, 39% carbohydrates). All animal experimentation was conducted in male mice on an inbred C57Bl6/J background following approval by local ethics committees (X9017/17). Unless stated otherwise, the animals were 12 to 16 weeks of age.

#### Quantitative protein analysis by SDS-PAGE and immunohistochemistry

Protein lysates for immunoblotting were generated by homogenization of human or murine tissues in RIPA buffer (50 mM Tris-HCl pH 7.4, 150 mM NaCl, 0.1% SDS, 0.5% sodium deoxycholate, 1% NP-40) containing complete Protease Inhibitor Cocktail (Sigma, 4693116001). Protein concentrations were determined using the BCA assay (Pierce, 23227). Lysates were adjusted for protein concentration and characterized by western blotting using standard SDS-PAGE. The intensities of immunoreactive signals were quantified by densitometric scanning of blots using the Image Studio Lite software. Primary antibodies used for immunodetection were directed against β-tubulin (abcam, ab6046), GAPDH (Genetex, GTX627408-01), apoE (Millipore, AB947), or sortilin (BD Transduction Laboratories™, 612101). Levels of APP and soluble APP processing products Aβ40, Aβ42, and sAPPβ in murine brain extracts were analyzed by commercial ELISA kits (Mesoscale Discovery).

For immunohistochemical detection of GFAP, 25 μm coronal cryosections of brain tissue were fixed with PFA, blocked in 1% horse serum in PBS, and incubated overnight at 4°C with Cy3-labeled anti-GFAP (Sigma, #C9205; 1:1,000) and biotinylated anti-NeuN (Chemicon, #MAB377B; 1:100) antibodies diluted in PBS (with 1% bovine serum albumin, 1% donkey serum, 0.3% TritonX-100). After washing with PBS, the sections were incubated with fluorescently labeled streptavidin diluted in 0.3% TritonX-100/PBS. Images were collected on a fluorescent microscope and analyzed using ImageJ. The area covered by GFAP immunosignals was quantified using the 'Analyze Particles' function after manual selection of cortical area and thresholding.

**Analysis of gene transcription**

Total RNA was extracted from tissue and cell lysates using the TRIzol reagent and purified with RNeasy Mini/Micro Kit (Qiagen). Reversely transcribed cDNA was generated from the total RNA using the RT2 First Strand Kit (Qiagen) and subjected to quantitative RT-PCR using Taqman or sybergreeen probes *Actb* (Mm02619580_g1), *Gapdh* (Mm99999915_g1), *B2m* (Mm00437762_m1), *Gfap* (Mm01253033_m1), *Tnfa* (Mm00443260_g1), *Vegf*

(Mm00437306_m1), *Sort1* (Mm00490905_m1), human *APOE* (Hs00171168_m1), *Mgll* (Mm00449274_m1), *Dagla* (Mm00813830_m1), *Faah* (Mm00515684_m1), *Napepld* (Mm00724596_m1), *Klf10* (PPM05167F-200), *Mmp9* (PPM03661C-200) and *Pparg* (PPM05108C-200). Fold change in transcript levels was calculated using the cycle threshold (CT) comparative method ($2^{-ddCT}$) normalizing to CT values of internal control genes *Gapdh* or *B2m* [4]. RT-PCR analysis of PPAR target genes was performed using the RT2 Profiler PCR Array in combination with RT2 SYBR green master mix according to the manufacturer's recommendations (PPAR Targets RT2 Profiler PCR Array, Qiagen). The comparison in transcript levels of different genes was calculated using the cycle threshold (CT) comparative method ($2^{-dCT}$) normalizing to CT values of internal control gene *Gapdh*.

**Lipid analyses**

For fatty acid profiling, 30 mg of human or mouse brain cortex were hydrolyzed with 100 µl 10 mol/l NaOH within 60 min at 80°C. The samples were neutralized with 100 µl acetic acid. A 50 µl aliquot was diluted 1:10 with methanol containing internal standards (50 µg of C15:0 and C21:0; 5 µg of C20:4-d8 and C18:2-d4, and 1 µg of C20:5-d5 and C22:6-d5; Cayman Chemical, Ann Arbor MI). HPLC measurements were performed using an Agilent 1290 HPLC system with binary pump, autosampler and column thermostat equipped with a Phenomenex Kinetex-C18 column 2.6 µm, 2.1 x 150 mm column (Phenomenex, Aschaffenburg) using a solvent system of acetic formic acid (0.1%) and acetonitrile. The solvent gradient started at 70 % acetonitrile and was increased to 98 % within 10 min and kept until 14 min with a flow rate of 0.4 ml/min and 5 µl injection volume. The HPLC was coupled with an Agilent 6470 triplequad mass spectrometer with electrospray ionisation source and operated in negative selected ion mode.

For eCB profiling, 10 mg cortex tissue were homogenized in 300 µl distilled water. Then, 25 µl citric acid (0.4 mol/l), 10 µl of internal standards (10 µg/ml DHA-Ethanolamid-

D4, Anandamid-D4, EPA-Ethanolamid-D4; Cayman Chemical, Ann Arbor MI) and 1 ml ethyl acetate were added. The samples were shaken for 10 min, centrifuged for 3 min at 3500 rpm and the upper phase recovered. Next, 1 ml ethyl acetate was added to the lower (sample) phase, followed by shaking for further 10 min and centrifugation for 3 min at 3500 rpm. Again, the upper phase was recovered and both supernatants combined. The solvent was removed to dryness at 40°C under a stream of $N_2$, before the pellets were resuspended in 100 µl acetonitrile and eCB measured. HPLC-measurement was performed using an Agilent 1290 HPLC system with binary pump, autosampler, and column thermostat equipped with a Phenomenex Kinetex-C18 column 2.6 µm, 2.1 x 150 mm column (Phenomenex, Aschaffenburg, DE) using a solvent system of formic acid (0.1%) and methanol. The solvent gradient started at 5% methanol and was increased to 95% within 10.6 min under gradient conditions and kept until 15 min with a flow rate of 0.7 ml/min and a 10 µl injection volume. The HPLC was coupled with an Agilent 6490 triplequad mass spectrometer with atmospheric pressure chemical ionization source operated in positive selected ion mode.

**Cell culture experiments**

To generate control cell medium or medium containing human apoE3 or apoE4, HEK293 cells were transiently transfected with expression constructs encoding for myc-tagged versions of human apoE3 or apoE4, or with the empty vector. Cell supernatants were conditioned for 48 hours in medium without FCS [5]. The concentration of apoE in media batches was determined by western blot analysis comparing apoE signal intensities to those from serial dilutions of recombinant human apoE (MBL, JM-4699-500) included in the same blots.

Primary glial cultures were prepared from newborn mice (postnatal day 0-1) using mechanical dissociation after enzymatic digestion with trypsin and DNAse. The tissue suspension was plated on poly-L-lysine coated flasks and maintained in DMEM supplemented with 10% FBS and penicillin/streptomycin (Invitrogen). After two days, the cultures were washed with PBS prior to medium change. Cells were cultured for 4 more days before samples were collected for Western blot and qRT-PCR analyses.

**Proximity ligation assays**

Proximity ligation is an experimental approach to test the spatial proximity of proteins in cells. Here, we employed oligonucleotide-conjugated secondary antibodies directed against two primary antisera recognizing sortilin or the myc-tag in apoE. Binding of both antibody species in close proximity allows for their hybridization by connector oligonucleotides, forming a circular DNA strand that can be amplified by PCR. Incorporation of fluorescence-labelled oligonucleotides in the PCR product enables localized detection of protein interaction. Here, Chinese hamster ovary cells stably overexpressing mouse sortilin (CHO-S) were grown on glass coverslips and treated for 24 hours with conditioned medium containing 5 μg/ml of human apoE3 or apoE4. Then, the cells were fixed with 4% PFA in PBS and subjected to proximity ligation assay (PLA) using Far-Red PLA probes, followed by ligation and amplification performed according to the manufacturer's recommendation (Sigma). Images were obtained on a SP5 confocal microscope.

**Trafficking studies in CHO cells**

To study sortilin recycling by immunocytochemistry (Fig. 6B-C), CHO-S cells were grown on glass coverslips. The cells were incubated with anti-sortilin antibodies (R&D Systems, AF2934) for 1 hour at 4°C to decorate sortilin molecules at the cell surface. To induce endocytosis of sortilin, cells were then placed for 15 minutes at 37°C in medium with 5 μg/ml of apoE3 or apoE4. Thereafter, the cells were incubated for another 60 minutes in medium containing 80 μM of dynasore (ab120192). Finally, the cells were fixed with 4% PFA in PBS and subjected to immunocytochemistry using Alexa Fluor 555-conjugated secondary antibody. Cells had been non-permeabilized or permeabilized with 0.3% of triton in PBS prior to incubation with secondary antiserum to distinguish between cell surface receptor and total receptor signals, respectively. Cells were mounted with DAKO fluorescence mounting medium.

To quantify the amount of sortilin molecules retained in cells in the presence of apoE3 or apoE4, CHO-S cells were grown to confluency in 6-well plates. Then, the cells were incubated with 0.5 mg/ml EZ-Link™ Sulfo-NHS-SS-Biotin (Pierce, 21331) for 30 minutes at 4°C to biotinylate surface proteins. Thereafter, the cells were placed at 37°C in medium containing 5 μg/ml of apoE3 or apoE4 to induce internalization of sortilin/apoE complexes. After 15 minutes, proteins retained at the cell surface were stripped of the biotin label using 50 mM glutathione (GSH; Sigma, G6529 in) in medium containing 75 mM sodium chloride, 1 mM magnesium chloride, 0.1 mM calcium chloride, 80 mM sodium hydroxide, 10% fetal bovine serum). Finally, the cells were incubated for another 60 minutes at 37°C in medium containing 80 μM of dynasore and GSH solution to remove biotin from all receptor molecules recycled back to the cell surface. The cells were scraped in lysis buffer (50 mM Tris HCL, pH 7.4, 140 mM NaCl and 1% Triton X-100) and cell lysates used for pull-down of biotinylated proteins on Streptavidin magnetic beads (Thermo Fisher 10150874). Beads were boiled in Laemmli buffer to release captured proteins and subjected to western blot analysis of sortilin.

**Statistical analysis**

For all *in vivo* experiments, an indicated number *n* is the number of mice per group used in an experiment. For primary culture experiments, an indicated number *n* is the number of independent glial preparations (biological replicates) used for western blotting or qRT-PCR analyses. For co-localization studies in CHO cells, *n* is the number of cells analyzed in replicate experiments. Each mouse (or biological replicate in a cell culture experiment) represents a statistically independent experimental unit, which was treated accordingly as an independent value in the statistical analyses. Statistical analyses were performed using GraphPad Prism software. For all other data with two independent variables (factors), two-way ANOVA with Bonferroni multiple comparisons test was applied. Further details of statistical analyses not included here are specified in the respective figure legends.

### SUPPLEMENTAL REFERENCES


[1] Games D, Adams D, Alessandrini R, Barbour R, Berthelette P, Blackwell C, et al. Alzheimer-type neuropathology in transgenic mice overexpressing V717F beta-amyloid precursor protein. Nature. 1995;373:523-7.

[2] Jansen P, Giehl K, Nyengaard JR, Teng K, Lioubinski O, Sjoegaard SS, et al. Roles for the pro-neurotrophin receptor sortilin in neuronal development, aging and brain injury. Nat Neurosci. 2007;10:1449-57.

[3] Knouff C, Hinsdale ME, Mezdour H, Altenburg MK, Watanabe M, Quarfordt SH, et al. Apo E structure determines VLDL clearance and atherosclerosis risk in mice. J Clin Invest. 1999;103:1579-86.

[4] Schmittgen TD, Livak KJ. Analyzing real-time PCR data by the comparative C(T) method. Nat Protoc. 2008;3:1101-8.

[5] Chen Y, Durakoglugil MS, Xian X, Herz J. ApoE4 reduces glutamate receptor function and synaptic plasticity by selectively impairing ApoE receptor recycling. Proc Natl Acad Sci U S A. 2010;107:12011-6.

### SUPPLEMENTAL FIGURE LEGENDS

**Figure S1. Sortilin deficiency does not impact apoE production in primary astrocytes**

**(A)** Western blot analysis of apoE3 and apoE4 levels in primary astrocytes from wild-type (WT) and sortilin-deficient (KO) mice expressing human apoE3 or apoE4. Detection of sortilin served as control. No difference in apoE levels was seen comparing WT and KO astrocytes on either apoE background. (**B**) Quantitative RT-PCR analysis of transcript levels for human apoE3 and 4 and sortilin (*Sort1*) in primary astrocytes of WT and KO mice (n=5 independent cell preparations per genotype). Values are mean ± SEM given as $\log_2$ fold change compared with levels in the respective WT sample (mean set to 0). Transcript levels for *Sort1* but not for *APOE* are impacted by the *Sort1* genotype (Two-way ANOVA, followed by Bonferroni post-hoc analysis). **, $p<0.01$; ***, $p<0.001$.

**Figure S2. Interaction of sortilin and apoE3 controls Aβ42 levels in hippocampus**

Levels of full-length APP (**A**), soluble Aβ40 (**B**), soluble Aβ42 (**C**), and sAPPβ (**D**) were determined by ELISA in hippocampal brain extracts of apoE3 or apoE4 targeted replacement mice crossed with the PDAPP strain (4 months of age). In addition, the animals were either wild-type (WT) or homozygous for the *Sort1* null allele (KO). Values are mean ± SEM (n = 8-12 animals per group). Significant differences in levels of APP and processing products were determined by Two-way ANOVA, followed by Bonferroni post-hoc analysis (**, $p<0.01$; ***, $p<0.001$)

**Figure S3. Levels of arachidonic acid and DHA in murine and human brains**

(**A**) Levels of arachidonic acid and DHA in brain cortices of apoE3- and apoE4-targeted replacement mice either wild-type (WT) or homozygous for the *Sort1* null allele (KO) (3 months of age, n=12 per genotype). Data are the mean ± SEM. The significance of data was determined by Two-way ANOVA, followed by Bonferroni post-hoc analysis (*, $p<0.05$; **, $p<0.01$; ***, $p<0.001$). (**B**) Levels of arachidonic acid and DHA in prefrontal cortex specimens of AD patients homozygous for *APOE3* or *APOE4* (n=10 for *APOE4*, n=22 for *APOE3*.) Values are mean ± SEM. Significance of data was determined by unpaired Student *t* test.

**Figure S4. Levels of arachidonic acid and DHA in plasma and brain lipoproteins**

(**A**) Plasma lipoprotein profiles of WT and *Sort1* KO mice expressing apoE3 (left panel) or apoE4 (right panel). Values are mean ± SEM (n = 3-6 animals per group). HDL, high density lipoprotein; LDL, low density lipoprotein; VLDL, very low density lipoprotein. (**B**) ApoE-containing HDL particles were purified from the plasma of mice of the indicated genotypes and analyzed for levels of arachidonic acid and DHA (n=3 per genotype group). No significant differences in lipoprotein profiles or DHA and arachidonic acid levels were detected comparing genotype groups. (**C**) ApoE-containing lipoproteins were purified from human cerebrospinal fluid (CSF) using heparin affinity chromatography. Enrichment of apoE in two eluate fractions (apoE) but not in the heparin column flow-through (non-apoE) is documented by western blotting. (**D-F**) Levels of total fatty acids (D), arachidonic acid (E), and DHA (F) in apoE-containing lipoproteins from CSF samples of AD patients (AD) and matched control subjects (Ctrl) are given. Donors were homozygous for *APOE3* or *APOE4*. No statistically significant differences in lipid levels were seen comparing the indicated groups (n=7-10 subjects per group).

**Figure S5. Brain transcript levels of enzymes in eCB metabolism**

Transcript levels of the indicated genes in total brain extracts from apoE3 and apoE4 mice, either WT or KO for *Sort1* were determined by quantitative RT-PCR. Values are mean ± SEM of $log_2$ fold change compared to (E3;WT) set to value 0 (n =6-9 mice per genotype group; Two-way ANOVA). Dagla, diacylglycerol lipase α; Faah, fatty acid amide hydrolase; Nape-pld, N-acyl phospathidylethanolamine phospholipase D; Mgll, monoglyceride lipase.

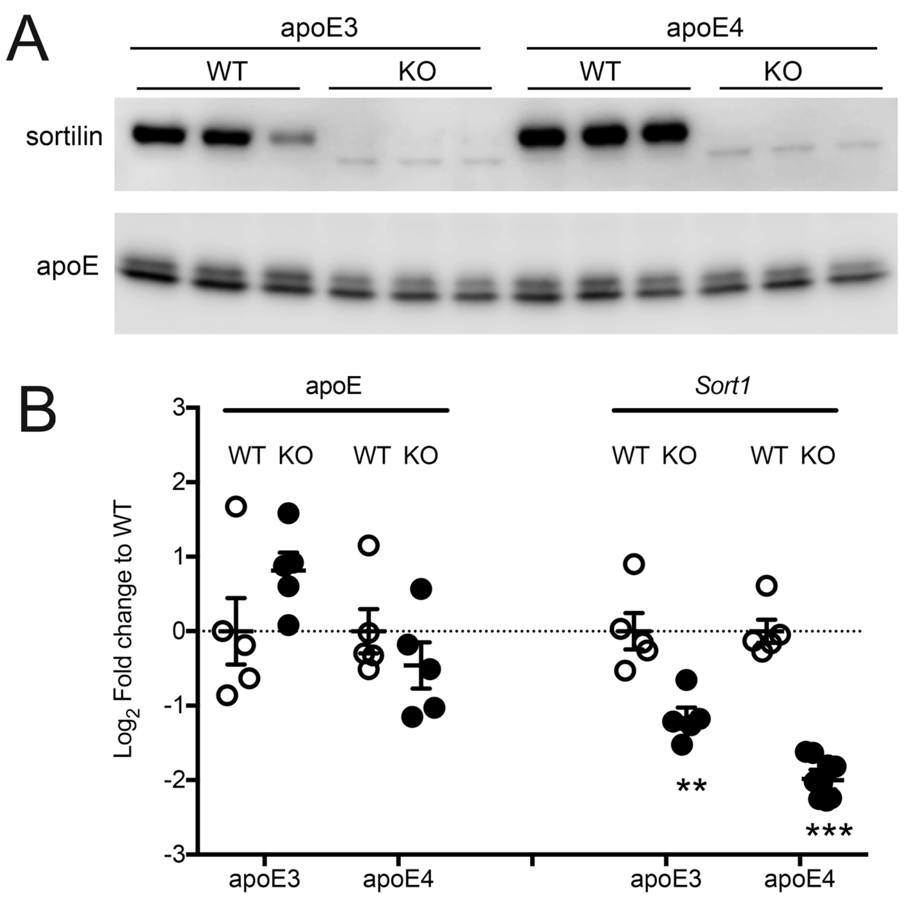


Asaro et al., Suppl. figure S1

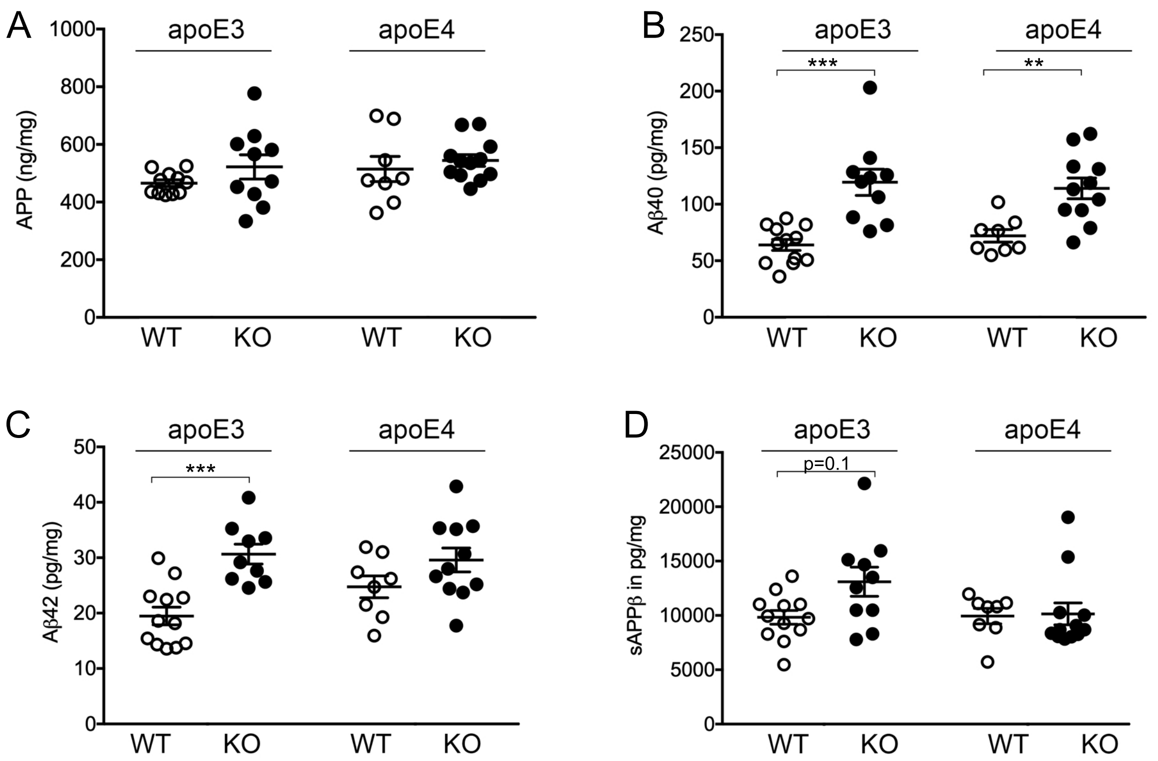


Asaro et al., Suppl. figure S2

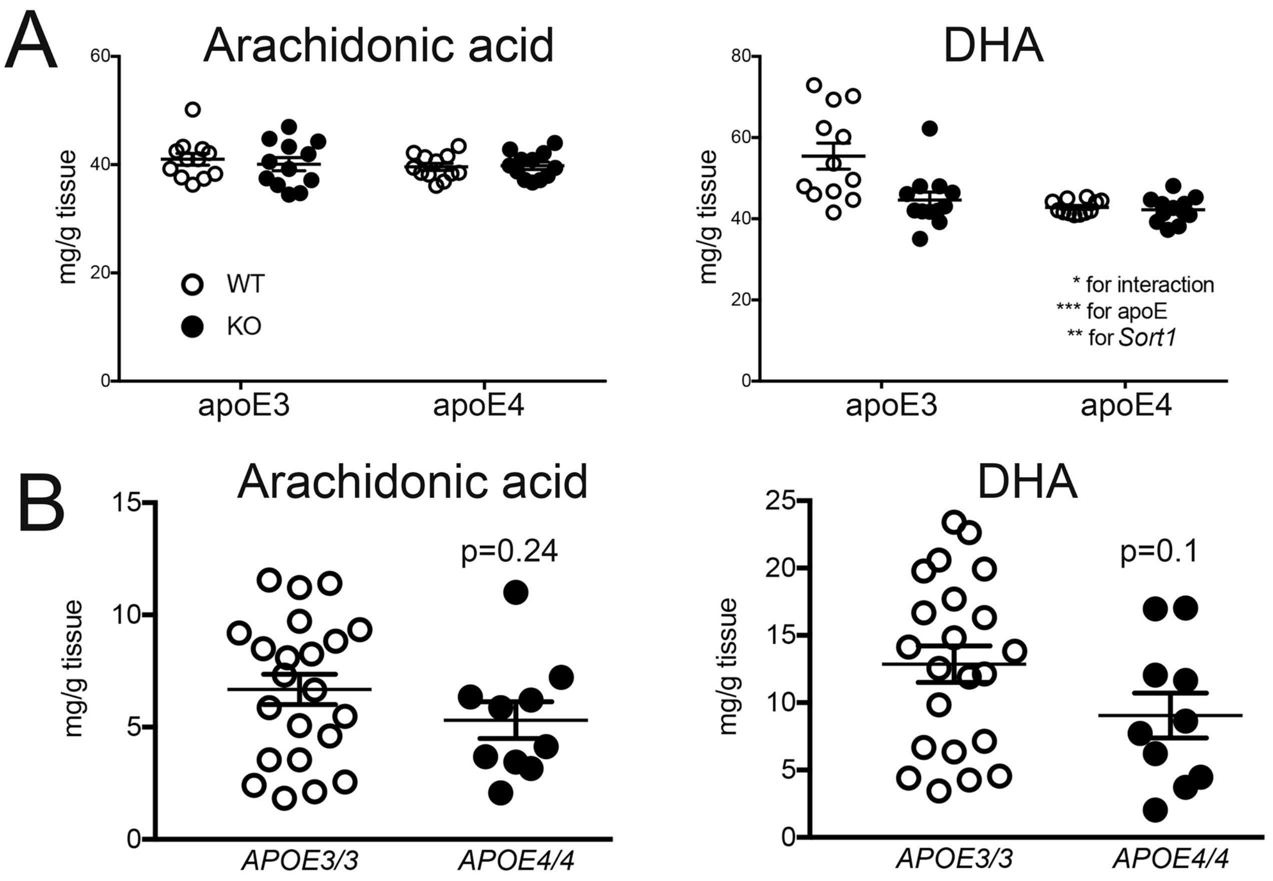


Asaro et al., Suppl. figure S3

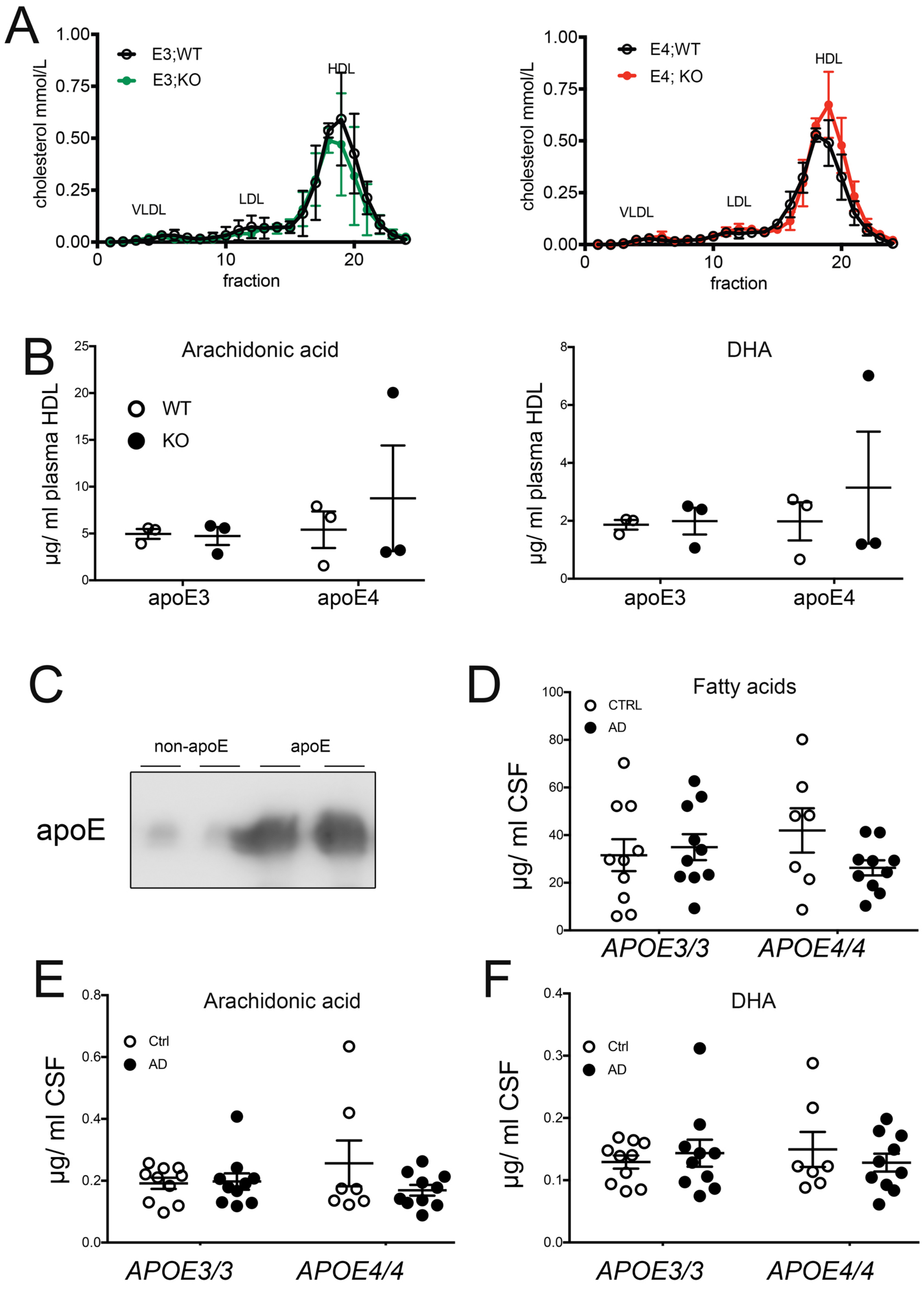


Asaro et al., Suppl. figure S4

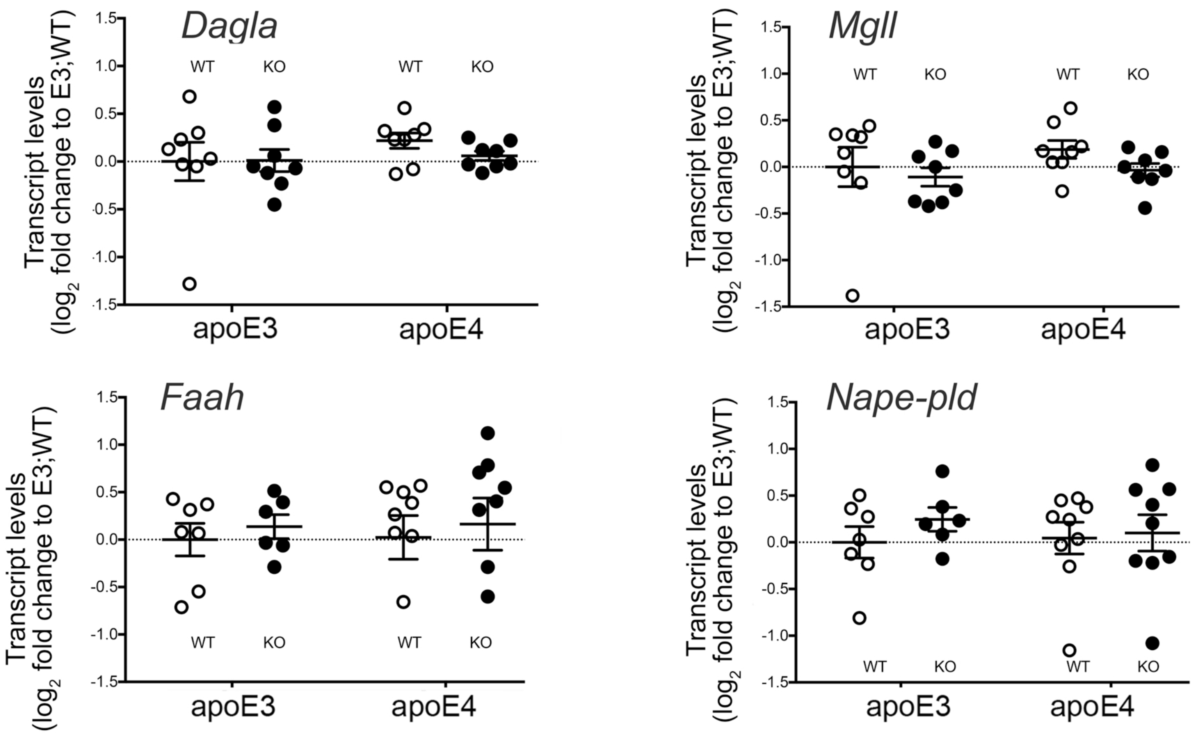


Asaro et al., Suppl. figure S5
